# Effect of vertical slit turbulence on metabolism and swimming behavior of juvenile grass carp (*ctenopharyngodon idella*)

**DOI:** 10.1101/190272

**Authors:** Yuan Xi, Jiang Qing, Xu Meng, Tu Zhi-ying, Zhou Yi-hong, Huang Ying-ping

## Abstract

Baffles were incorporated into the swim chamber of a flume-type swimming respirometer, and the effect of vertical slit turbulence on the swimming behavior and metabolism of juvenile grass carp were investigated. Results showed a significant lower *TBF* in turbulent flow than in laminar flow (*p*< 0.05). However, differences in *TBF* at different inlet velocities were not significant (*p*> 0.05), whether the fish passed through the baffles or not. In turbulent flow, the residence time ratios of test fish at different flow zone were low water velocity > medium velocity > high velocity. Oxygen consumption rate (*MO*_2_) increased with flow velocity and was higher in turbulent flow than in laminar flow. Further, the speed exponent c, in turbulent flow, was significantly higher than in laminar flow, indicated a decrease swimming efficiency. This study of fish swimming in turbulent flow extends knowledge of fish ecology and provides data for guiding the design of hydrokinetic turbulent where needed, so preventing ecological impacts.

## Introduction

The ability of fish to stabilize body posture and position, and swimming strategies to minimize potential negative effects or maximize potential benefits differ with flow rate and pattern. Fish swimming behavior and energy consumption varied with habitat and activity; feeding, migration, schooling, etc. (Yan et al., 2015). For example, oxygen consumption increases with swimming speed at optimal water temperatures (Claireaux et al., 2006; Fuiman & Batty, 1997; Lee et al., 2003). Present fish swimming behavior studies are generally conducted in near laminar flow, but uniform, steady flow is unusual in natural habitats. Swimming behavior and metabolism of fish obtained in laminar flow can’t reflect natural conditions, and the confined space of a respirometer swimming chamber limits swimming performance. Thus, engineering applications (e.g. fishway design) of fish swim test data, obtained under controlled conditions, has been questioned.

The effects of turbulence and variable flow on fish behavior has not been well studied, in part because turbulent flow, by definition, is complex. The strength and range dynamics of turbulence can attract or repel and promote or hinder movement during migration. Some fish species are able to take advantage of columnar vortices to decrease metabolic cost by entrainment and Kármán gaiting (Liao, 2007; Przybilla et al., 2010; Taguchi & Liao, 2011). These studies have focused mostly on vortices produced by stationary semicylinders, positioned horizontally or vertically (Liao, 2007; Taguchi & Liao, 2011; Tritico & Cotel, 2010). Several groups have found increased fish abundance in areas of streams with high turbulence (Smith et al., 2005). However, turbulence produced by horizontal cylinders leads to avoidance behavior (Eidietis et al., 2002; Webb, 1998; Webb & Cotel, 2010). Finally, other studies have found that turbulence generally increases energy consumption (Enders et al., 2003; Lupandin, 2005; Roche et al., 2014; Tritico & Cotel, 2010) and small turbulent eddies may interfere with sensory receptors (Webb & Cotel, 2010). Thus, the study of fish swimming ability and behavior in turbulent flow will extend our knowledge of fish ecology and has significance for engineering applications.

Grass carp (*ctenopharyngodon idellus*) is a migratory fish inhabiting rivers and lakes, and is a target species for many fish facilities. Information on swimming behavior in non-uniform flow fields is urgently needed. In this investigation, the swimming ability and behavior of juvenile grass carp were studied in turbulent flow, simulated using FLOW-3D software. A flume-type respirometer was fitted with two baffles of unequal width to produce slit-baffle turbulence, and fish metabolism (oxygen consumption rate) and behavior (tail beat frequency, *TBF*) were investigated; and the time spent in different velocity regions. The results provide a reference for fish adaptability to different river hydraulic conditions.

## Methods

### Test fish and acclimation procedure

Juvenile grass carp obtained from a fish hatchery in Yichang. Sixty fish (body length, 8.0-10.0 cm; body mass, 9.2-13.0 g) were maintained in a recirculating water tank system for two weeks with a natural photoperiod in fresh dechlorinated water at natural temperature (10 ± 0.50°C), *DO* (dissolved oxygen) > 7.00 mg L^-1^, pH 6.50 −7.30, ammonia-N 0.005-0.025 ppm. Fish were fed daily to satiation at 9:00 am with a commercial diet and residual food and feces were removed. All measurements were conducted after 48 h of fasting to ensure a post-absorptive state. Twelve carp were tested in each trial. Fish were individually transferred into the swim chamber and allowed to acclimate at a low current velocity (0.02 m s^-1^) for 2 h. The water temperature in the swimming chamber was controlled within ±1.0°C. The velocity was increased in 1.0 BL increments each 30 min until the fish was exhausted (failure to move away from the swimming chamber screen for 20 s), or the test fish was past the slit for ~ 20 s. After each velocity increment, the respirometer was flushed with aerated water from an external tank for 5 min to keep dissolved oxygen levels above 80% saturation.

### Experimental apparatus

Tests were carried out in a temperature controlled flume-type respirometer (Fig 1) with a 6 L swimming channel, 60 cm × 10 cm × 10 cm (L × W × H) and sensor mounts for temperature and *DO*. The flow rate was measured with a current meter (Vectrino, Nortek) and *DO* was measured using a dissolved oxygen meter (Hach, USA, range, 0 - 20 mg L^-1^, precision, 0.01 mg L^-1^). The flow rate was controlled with a propeller driven by a 370 watt variable-speed motor. The recirculating water was passed through a honeycomb (1 cm cell diameter), placed upstream of the chamber, to produce near-laminar flow at the swim chamber entrance. The water velocity was range of 0.02-1.37 m s^-1^. To record swimming behavior and determine residence time and tail beat frequency (TBF), a CCD video camera was placed directly over and perpendicular to the swim chamber at a height of 0.50 m. Two baffles of 1 cm and 2 cm were used to form a vertical slit, producing turbulent flow in the swim chamber (Fig 2).

**Fig 1.**
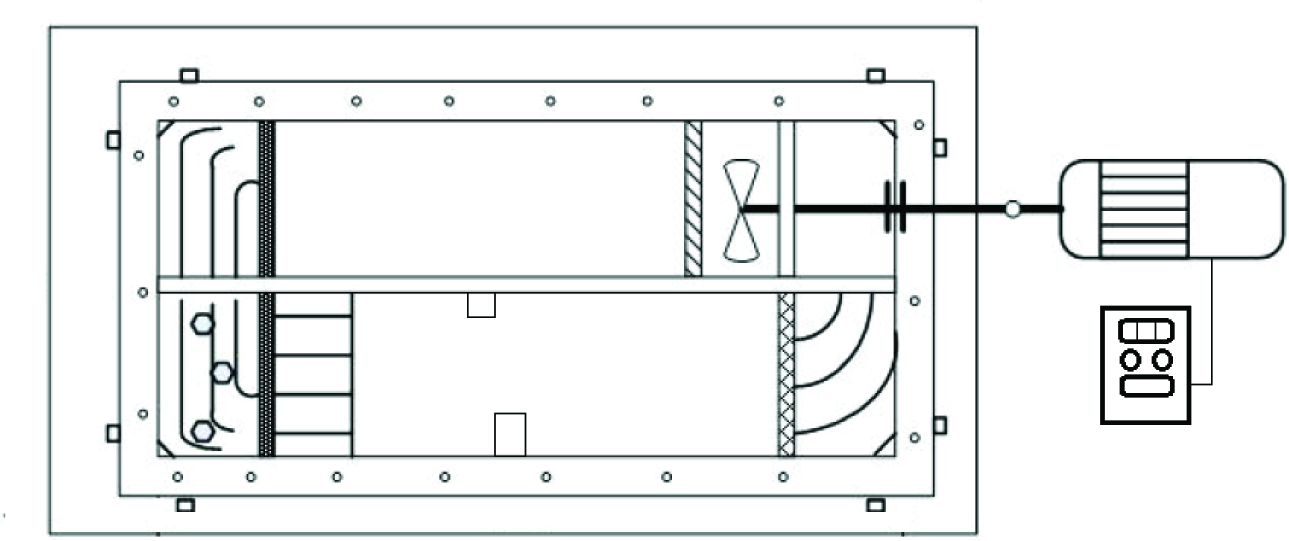
Experiment device for fish swimming capability test. Two baffles of 1 cm and 2 cm were set to form a vertical slit, producing turbulent flow in the swim chamber; and laminar flow without baffles in the swimming zone.

**Fig 2.**
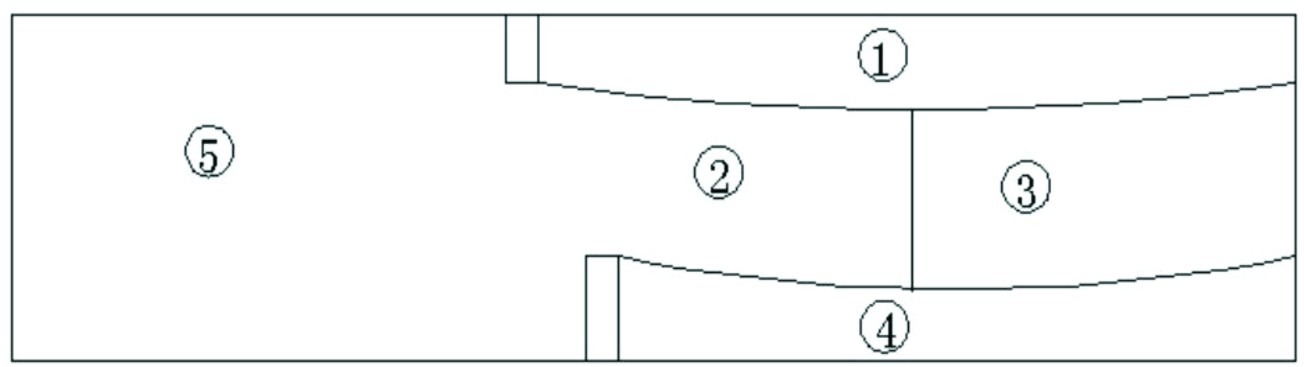
Different velocity distribution in experimental area; the flow velocity zone below the slit was divided to four flow velocity zones; (1) behind the short baffle and (4) behind the long baffle were low velocity zones, (2) was the high water velocity zone, and (3) was the medium water velocity zon, and (5) was above the slit.

Turbulence in the swim chamber was modeled with a three-dimensional finite element flow model, Flow-3D, using renormalization of turbulence energy and dissipation rate. The model was calibrated using detailed measurement of 3D flow velocities, obtained with the current meter. The model was then applied to predict flow velocity and maximum turbulent kinetic energy (TKE_max_) around the vertical slit. The relationship between the inlet water velocity (*U*) and the maximum water velocity (U_max_) and TKE_max_ were; U_max_=1.38*U* (*R*^2^=1) (Fig 3) and TKE_max_=0.092*U*^0.62^ (*R*^2^=1) (Fig 4), respectively.

**Fig 3.**
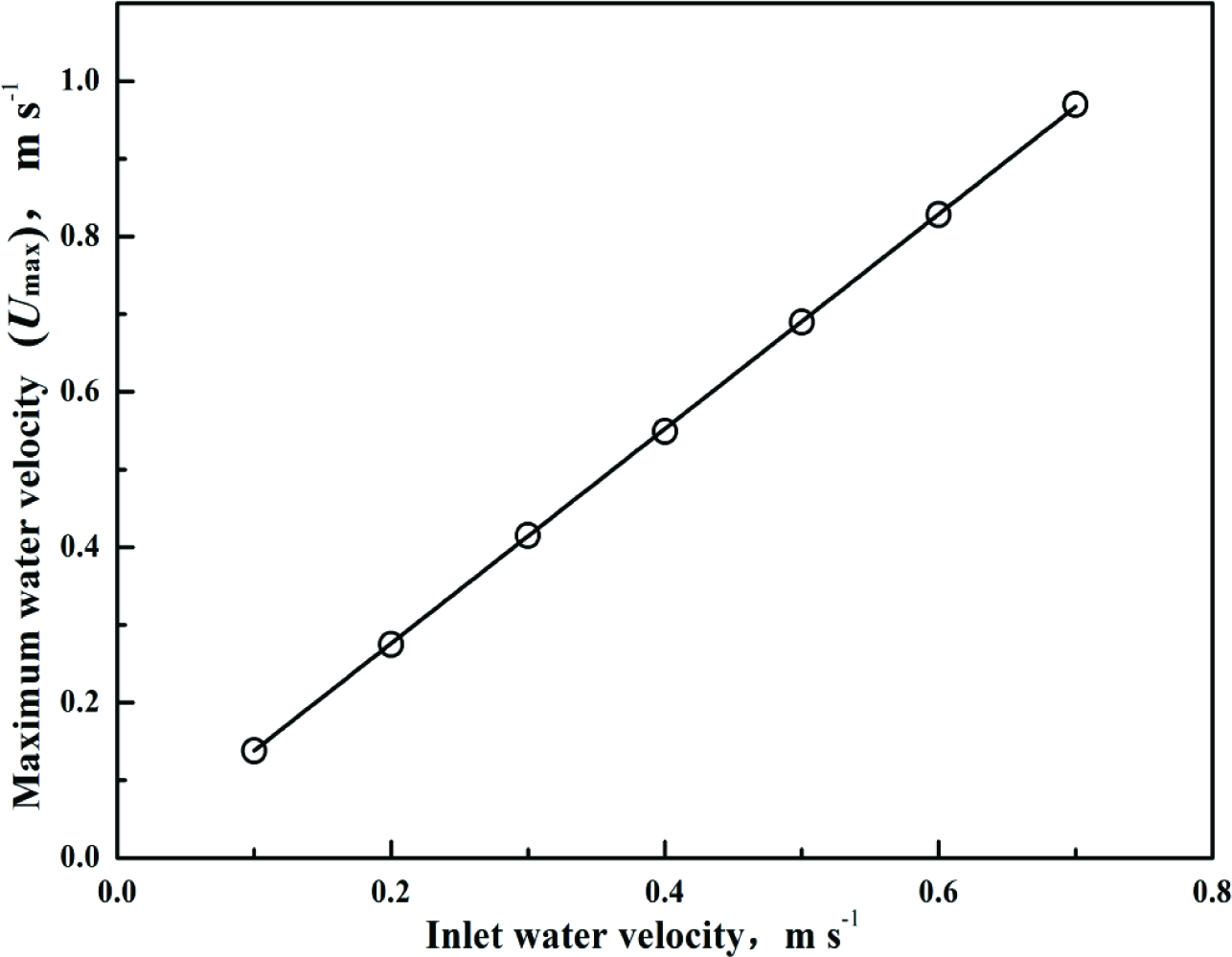
Relationships between inlet water velocity and maximum water velocity (*U*_max_), *U*_max_=1.38U (*R*^2^=1).

**Fig 4.**
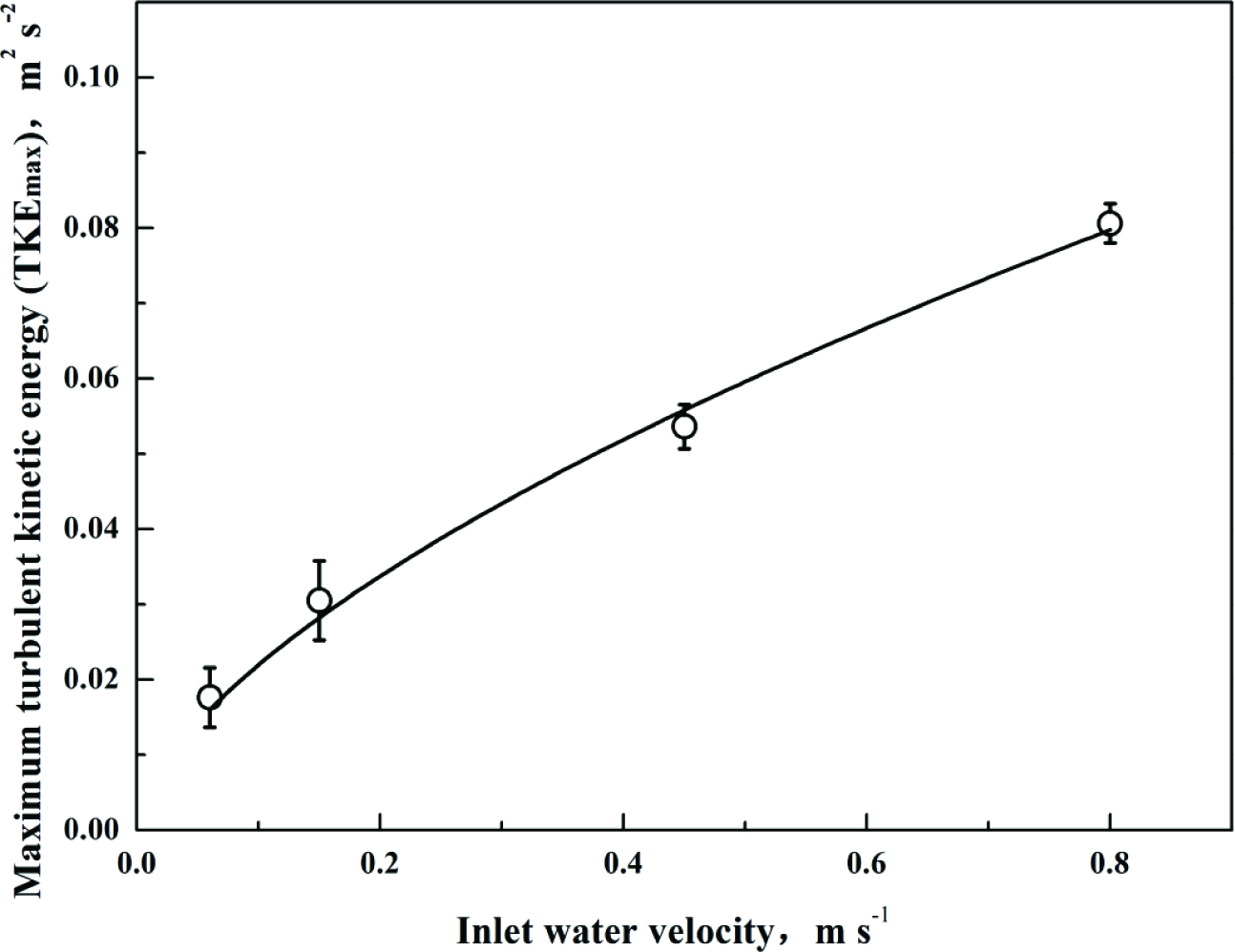
Relationships between inlet water velocity and maximum turbulent kinetic energy (TKE_max_), TKE_max_=0.092*U*^0.62^ (*R*^2^=1). In this study, the inlet velocity range was 1-4 BL/s, resulting in a turbulent kinetic energy range of 0.022-0.052 m^2^ s^-2^.

### Oxygen consumption rate

The weight-specific oxygen consumption rate (*MO*_2_, mgO_2_ kg^-1^ h^-1^) for each velocity increment was determined by recording the decrease in *DO* every 5 min in the sealed respirometer for the middle 20 min of the 30 min time step. *MO*_2_ was calculated using Equation 1 with the slope of the regression line obtained by plotting *DO* against time:

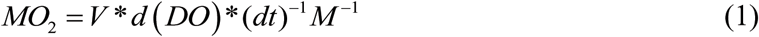

where *V* is the respirometer volume (L); *d*(*DO*)/*dt*, the slope (mg O_2_ h^-1^); and *M* the fish mass (kg). The rate was corrected for background *DO* consumption by measuring *DO* in the absence of fish.

Linear, exponential, and power functions were used to describe the relationship between oxygen consumption rate and swimming speed (Webb, 1993) and the power function gave highest regression coefficients and Equation 2 was used to describe the *MO*_2_-*U* relationship:

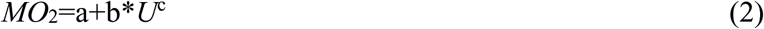

Where a, b, and c are constants; and c is the speed exponent.

### Residence time ratio and *TBF*

Grass carp swimming was tracked at 25 frames per second during each time step with a CCD-camera mounted at a height of 0.5 m above the respirometer. The first 10 min were excluded from the analysis to avoid the initial response to change in flow and each swimming behavior was established during a period of 5 min. Video recordings were analyzed to obtain residence time in each flow zone (the entire head and pectoral fins had to be in the zone to be counted), and the ratio was calculated. The same video recordings were used to determine *TBF* by counting the number of complete tail beat cycles per minute.

### Data analyses

Statistical comparisons among all fish were accomplished using a parametric analysis of variance (ANOVA). In cases where ANOVA detected significant differences, a pairwise post-hoc Duncan test was used to determine difference. Values are reported as mean ± S.E.M and differences were considered significant at *p*< 0.05.

## Results

### Residence time ratio in the flow rate regions

Analysis of fish motion was used to obtain the residence time ratio in different turbulent flow velocity zones (Fig 4). The residence time ratios followed the order of velocity; (4) flow velocity zone> (1) flow velocity zone> (3) flow velocity zone> (2) flow velocity zone. Residence time was significantly longer in the low flow velocity zones than in the high flow velocity zone (*p*< 0.05). The difference in residence times was not significant, but time behind the long baffle was longer than time behind the short baffle (*p*> 0.05). At the low inlet flow velocity, there was no significant difference in residence time between flow velocity Zone (2) and Zone (3). Residence time in flow velocity Zone (2) increased with inlet flow velocity and was significantly higher at a flow velocity of 4 BL s^-1^ (*p*< 0.05), and the sprint frequency to pass the baffle increased.

### TBF

At an inlet water velocity of 4 BL s^-1^, juvenile grass carp either passed through the vertical slot or failed to continue swimming. *TBF* increased linearly with swimming speed (Fig 5) up to *U*=4 BL s^-1^, when the fish either passed through the baffles or failed to resume swimming. Among the twelve juvenile grass carp tested, eight passed through the baffle, and four did not. The slope (TB/BL), intercept and correlation coefficient at each test trail are given in Table 1. The reciprocal of the slope (*BL/TB*) gives the stride length (*L*_*s*_). In turbulent flow, differences in *TBF* at different inlet velocities were not significant (*p*> 0.05), whether the fish passed through the baffles or not. However, *TBF* was significant higher in turbulent flow than in laminar flow (*p*< 0.05).

**Fig 5.**
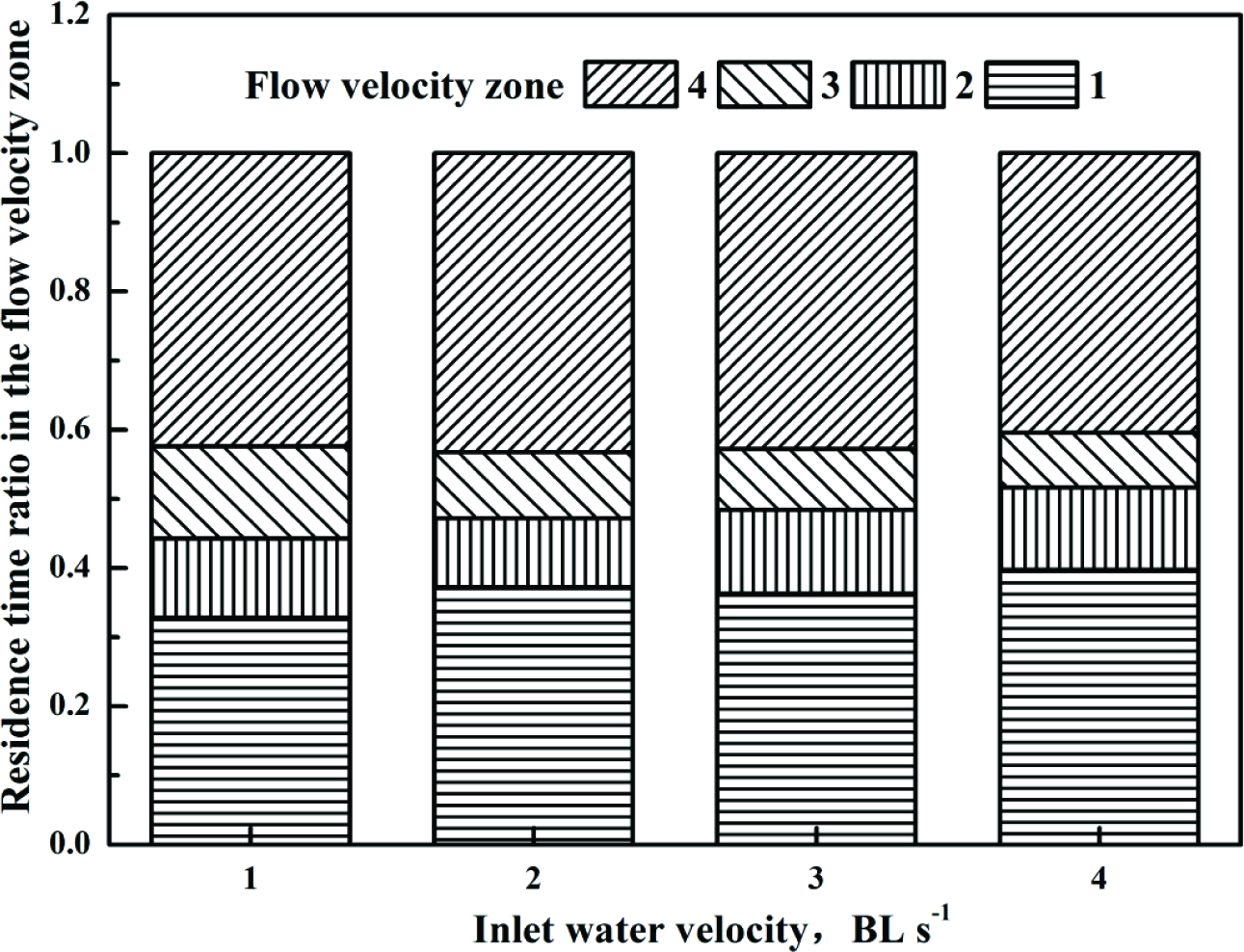
Residence time ratio of juvenile grass carp in various flow velocity zones at different inlet water velocities.

**Table. 1.** Intercept (a) and slope (b) for turbulent flow and laminar flow groups from the regression ***TBF*** =*a*+*b*\****U***, with fit indicated by *R*^2^.

| trail | Laminar flow | Pass baffles in<br>turbulent flow | Not pass baffles in<br>turbulent flow |
| --- | --- | --- | --- |
| $a$ | 1.57 | -0.08 | -0.04 |
| $b$ | 0.52 | 0.74 | 0.74 |
| $R^2$ | 0.98 | 0.94 | 0.99 |

### Oxygen consumption rate

Mean oxygen consumption rate (*MO*_2_) increased significantly with swimming speed *U* (BL s^-1^) (Fig. 6) and the relationship was accurately described by a power function (Tables 2). Oxygen consumption rate (*MO*_2_) was higher in turbulent flow than in laminar flow. And the speed exponent c, in turbulent flow, was also significantly higher than in laminar flow.

**Fig 6.**
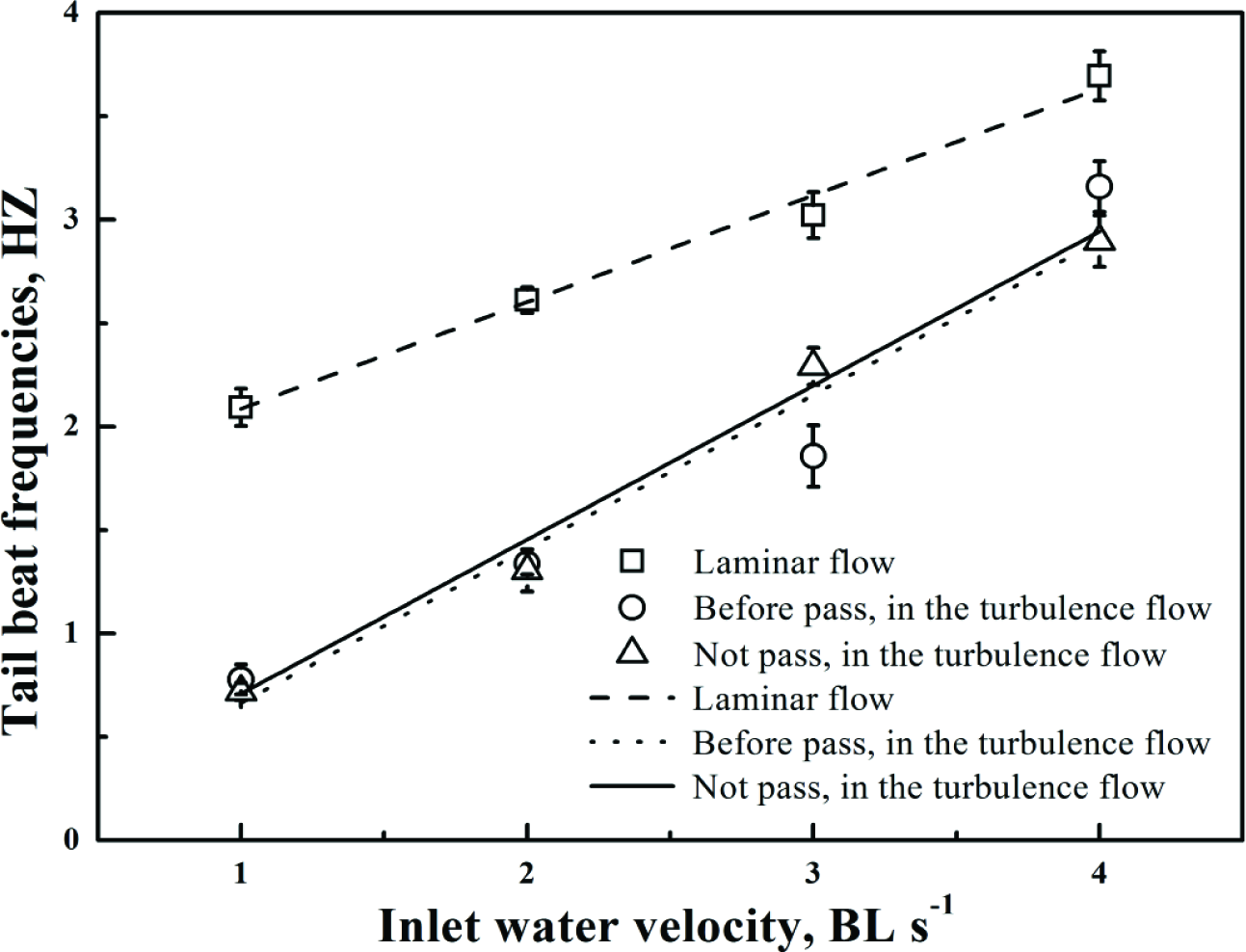
Relationship between inlet water velocity (*U*) and tail beat frequency (*TBF*) of juvenile grass carp in laminar flow and turbulent flow.

**Fig 7.**
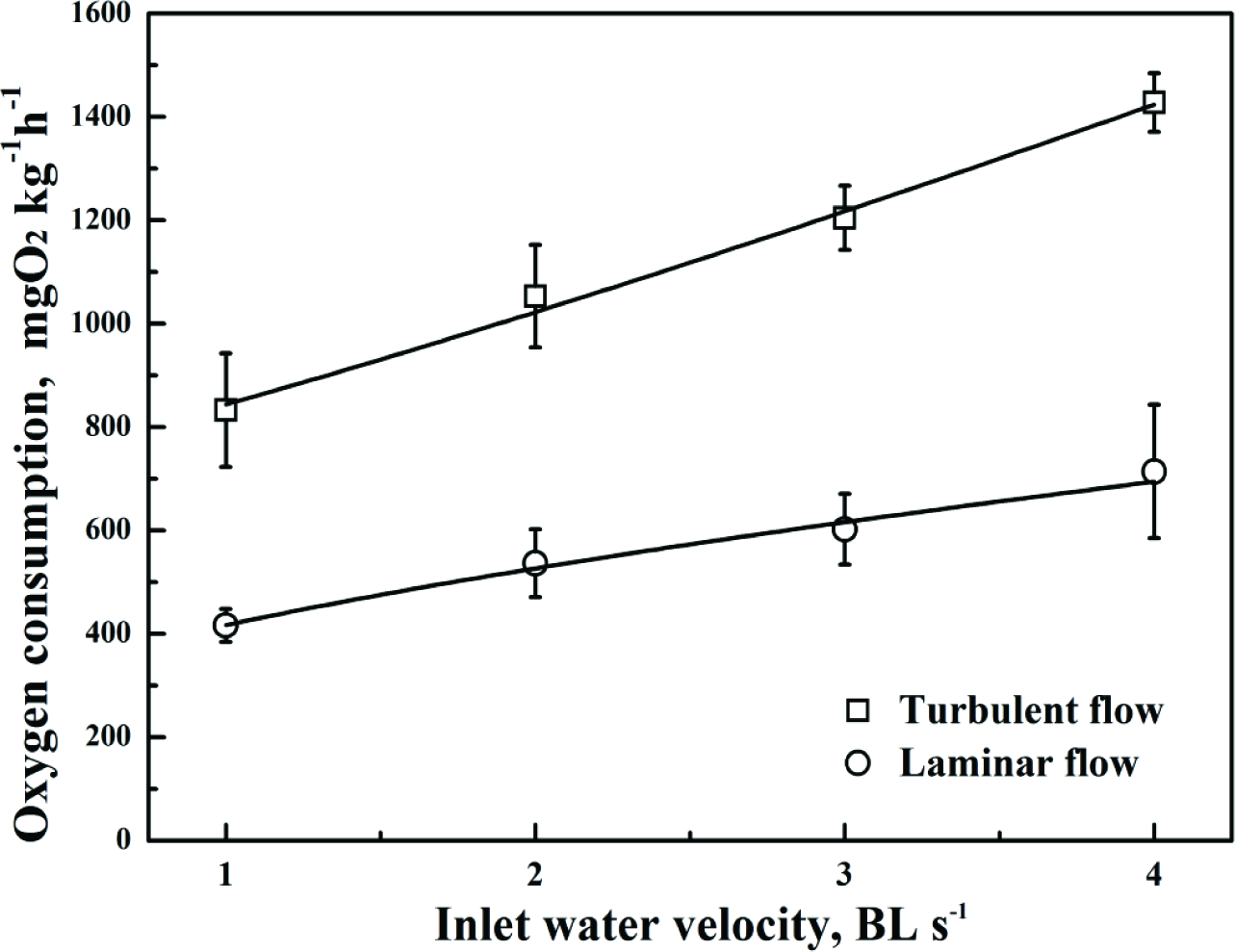
Relationship between inlet water velocity (*U*) and oxygen consumption (*MO*_2_) of juvenile grass carp in laminar flow and turbulent flow.

**Table. 2.** Coefficients (a, b, c) for turbulent flow and laminar flow groups of regression equation, *MO*_2_=*a*+*bU*^c^, with fit indicated by *R*^2^.

| trail | Turbulent flow | Laminar flow |
| --- | --- | --- |
| $a$ | 700.91 | 207.59 |
| $b$ | 142.94 | 209.28 |
| $c$ | 1.17 | 0.61 |
| $R^2$ | 0.98 | 0.98 |

## Discussion

In this study, the swimming behavior and oxygen consumption of juvenile grass carp was investigated under controlled conditions in laminar flow and turbulent flow.

Fish are subjected to different degrees of turbulence and may take advantage of the turbulence, depending on individual behavior and physiological state (Lupandin, 2005; Liao, 2007). Before passing through the baffles, residence time in the low velocity zones (1, 4) was significant longer than residence time in the high velocity zones (2, 3), indicating that juvenile grass carp prefer low flow velocity and low turbulence. In vertical slit fishways, there is a recirculation zone behind the baffle(s), with lower flow velocity and flow direction reversal, and it is a preferred location for fish (Rodriguez et al. 2006). When the inlet water velocity was low, the turbulent kinetic energy in Zone (2) was larger than in Zone (3). The avoidance behavior of juvenile grass carp was sufficient to result in a significantly shorter residence time ratio in Zone (2). As inlet velocity increased, the fish struggled to maintain balance and the stroke-glide ratio increased. As the fish attempted to pass through the vertical slot, the residence time ratio in Zone (2) increased significantly. It is consistent with the results of David et al. (2005) who found that juvenile *Oncorhynchus mykiss* preferred low turbulence zones at low flow velocity, but tended toward high turbulence zones at high flow velocity. In natural habitats, a high proportion of fish were observed in high turbulent zones (David et al., 2006).

Swimming costs was closely related to the ability of individual fish to adjust their fin kinematics to hold station in the swim chamber (Roche et al., 2014). The caudal fins of fish are remarkable propulsive devices and a key feature of fish evolution. *TBF* increases with swimming velocity, and the relationship can be linear or exponential (Ohlberger et al., 2007; Xiong & Lauder, 2014), and is used as an indicator of exercise performance (Herskin & Steffensen, 1999; Ohlberger et al., 2007). In this study, the relationship between *TBF* and swimming speed of juvenile grass carp was linear in both laminar and turbulent flow, but with significantly different *TBFs*. The slope (*b*=*TB BL*^-1^) was 0.74 in turbulent flow and 0.52 in laminar flow. It indicated that individuals with higher *TBF* were better at holding station and consumed less oxygen than fish with low *TBF*.

Further, the *TBF* of *Salmo playtcephalus* synchronized with the vortex frequency when adapting to turbulence behind a semicylinder (Liao et al., 2003). And swimming is discontinuous in turbulent flow, resulting in a significantly lower *TBF*, but a significantly higher *MO*_2_.

A change in swimming behavior will change metabolism and energy demand (Caputo et al., 2006), and cost of transport is a basic consideration for fish migration (Ohlberger et al. 2007). Swimming upstream in high turbulence can lead to disorientation and rapid increase in energy consumption (Enders et al., 2003; Silva et al., 2012) and may even result in damage to the fish (Cada et al., 1999; Odeh et al., 2002). Energy consumption in response to turbulence was ~14% of the total energy consumption in *Salmo salar* (Enders et al., 2005). The *MO*_2_ of juvenile grass carp was significantly higher in turbulent flow than in laminar flow. The value of c for turbulent flow was 1.17, it was significantly higher than in laminar flow (0.61) (Table 2). The value of c is inversely related to swimming efficiency (Tu et al., 2011). Thus, it decreases swimming efficiency of juvenile grass carp, in the turbulent kinetic energy range of 0.022-0.052 m^2^ s^-2^.

## Conclusion

Swimming behavior and metabolism of juvenile grass carp were affected by the turbulent field produced by baffles incorporated into the fish respirometer. In turbulent flow, *TBF* was significantly lower and *MO*_2_ significantly higher than in laminar flow. In turbulent flow, synchronization of swimming with vortex frequency resulted in discontinuous swimming and increased energy consumption. Turbulent flow has a complex vortex structure with vortex formations of different shape and size. In this study, the vertical slit turbulence intensity was relatively high for the body length (10 cm) of the test fish. In future research, we will test different configurations of baffle and swim chamber to determine if there are turbulence patterns and intensities that minimize energy consumption.

## Acknowledgments

This work was supported by Hubei Province Innovation Group Project (2015CFA021), the National Nature Science Foundation of China (51679126), and by the National Key Research Project (2016YFD0800902).

## Reference

Cada, G., Carlson, T., Ferguson, J., Richmond, M., Sale, M. (1999). Exploring the role of shear stress and severe turbulence in downstream fish passage. Office of Scientific & Technical Information Technical Reports, 1-9.

Caputo, F., Denadai, B. S., Greco, C. C. (2006). Intrinsic factors of the locomotion energy cost during swimming. Revista Brasileira De Medicina Do Esporte, 12, 399-404.

Cotel, J. A., Webb, W. P., Tritico, H. (2006). Do brown trout choose locations with reduced turbulence. Transactions of the American Fisheries Society, 135, 610-619.

David, L. S., Ernest, L. B., Mufeed, O. (2005). Response of juvenile rainbow trout to turbulence produced by prismatoidal shapes. Transactions of the American Fisheries Society, 134, 741-753.

David, L. S., Ernest, L. B., Bahman, S., Mufeed, O. (2006). Use of the average and fluctuating velocity components for estimation of volitional rainbow trout density. Transactions of the American Fisheries Society, 135, 431-441.

Enders, E. C., Boisclair, D., Roy, A. G. (2003). The effect of turbulence on the cost of swimming for juvenile Atlantic salmon. Canadian Journal of Fisheries & Aquatic Sciences, 60, 1149-1160.

Enders, E. C., Boisclair, D., Roy, A. G. (2005). A model of total swimming costs in turbulent flow for juvenile Atlantic salmon (Salmo salar). Canadian Journal of Fisheries & Aquatic Sciences, 62, 1079-1089.

Farrell, A. P. (2008). Comparisons of swimming performance in rainbow trout using constant acceleration and critical swimming speed tests. Journal of Fish Biology, 72, 693-710.

Freeman, M. C, Bowen, Z. H., Bovee, K. D., Irwin, E. R. (2008). Flow and habitat effects on juvenile fish abundance in natural and altered flow regimes. Ecological Applications, 11, 179-190.

Herskin, J & Steffensen. J. F. (1998). Energy savings in sea bass swimming in a school: measurements of tail beat frequency and oxygen consumption at different swimming speeds. Journal of Fish Biology, 53, 366-376.

Lea, J. M., Keen, A. N., Nudds, R. L., Shiels, H. A. (2016). Kinematics and energetics of swimming performance during acute warming in brown trout salmo trutta. Journal of Fish Biology, 88, 403-417.

Liao, J. C. (2007). A review of fish swimming mechanics and behaviour in altered flows. Philosophical Transactions of the Royal Society B Biological Sciences, 362, 1973-1993.

Liao, J. C, Beal, D. N., Lauder, G. V., Triantafyllou, M. S. (2003). Fish exploiting vortices decrease muscle activity. Science, 302, 1566-1569

Lupandin, A. I. (2005). Effect of flow turbulence on swimming speed of fish. Biology Bulletin, 32, 461-466.

Muller, U. K., Stamhuis, E. J., Videler, J. J. (2000). Hydrodynamics of unsteady fish swimming and the effects of body size: comparing the flow fields of fish larvae and adults. Journal of Experimental Biology, 203, 193-206.

Odeh, M. (2002). Evaluation of the effects of turbulence on the behavior of migratory fish, 2002 final report. Office of Scientific & Technical Information Technical Reports, 1-46.

Ohlberger, J., Staaks, G., Hölker, F. (2007). Estimating the active metabolic rate (AMR) in fish based on tail beat frequency (TBF) and body mass. Journal of Experimental Zoology Part A Ecological Genetics & Physiology, 307, 296-300.

Plaut, I. (2002). Critical swimming speed: its ecological relevance. Comparative Biochemistry & Physiology Part A Molecular & Integrative Physiology, 131, 41-50.

Roche, D. G., Taylor, M. K., Binning, S. A., Johansen, J. L., Domenici, P., Steffensen, J. F. (2014). Unsteady flow affects swimming energetics in a labriform fish (cymatogaster aggregata). Journal of Experimental Biology, 217, 414-422.

Rodriguez, T. T., Agudo, J. P., Mosquera, L. P., González, E. P. (2006). Evaluating vertical-slot fishway designs in terms of fish swimming capabilities. Ecological Engineering, 27, 37-48.

Silva, A. T., Katopodis, C., Santos, J. M., Ferreira, M. T., Pinheiro, A. N. (2012). Cyprinid swimming behaviour in response to turbulent flow. Ecological Engineering, 44, 314-328.

Swanson, C., Young, P. (1998). Swimming performance of delta smelt: maximum performance, and behavioral and kinematic limitations on swimming at submaximal velocities. Journal of Experimental Biology, 26, 333-45.

Tritico, H. M., Cotel, A. J. (2010). The effects of turbulent eddies on the stability and critical swimming speed of creek chub (semotilus atromaculatus). Journal of Experimental Biology, 213, 2284.

Tu, Z. Y., Yuan, X., Han, J. C., Shi, X. T., Huang, Y. P., David, M. J. (2011). Aerobic swimming performance of juvenile Schizothorax chongi (Pisces, Cyprinidae) in the Yalong River, southwestern China. Hydrobiologia, 675, 119-127.

Tudorache, C., Viaenen, P., Blust, R., Boeck, G. D. (2007). Longer flumes increase critical swimming speeds by increasing burst-glide swimming duration in carp cyprinus carpiom L. Journal of Fish Biology, 71, 1630-1638.

Xia, J., Ma, Y., Fu, C., Fu, S., Cooke, S. J. (2016). Effects of temperature acclimation on the critical thermal limits and swimming performance of brachymystax lenok tsinlingensis: a threatened fish in qinling mountain region of china. Ecological Research, 32, 61-70.

Xiong, G., Lauder, G. V. (2014). Center of mass motion in swimming fish: effects of speed and locomotor mode during undulatory propulsion. Zoology, 117, 269-281.

Yan, G. J., He, X. K., Cao, Z. D., Fu, S. J. (2015). Effects of fasting and feeding on the fast-start swimming performance of southern catfish Silurus meridionalis. Journal of Fish Biology, 86, 605-614.

